# Estimating the causal tissues for complex traits and diseases

**DOI:** 10.1101/074682

**Authors:** Halit Ongen, Andrew A. Brown, Olivier Delaneau, Nikolaos Panousis, Alexandra C. Nica, GTEx Consortium, Emmanouil T. Dermitzakis

## Abstract

Interpretation of biological causes of the predisposing markers identified through Genome Wide Association Studies (GWAS) remains an open question^1^. One direct and powerful way to assess the genetic causality behind GWAS is through expression quantitative trait loci (eQTLs)^2^. Here we describe a novel approach to estimate the tissues giving rise to the genetic causality behind a wide variety of GWAS traits, using the cis-eQTLs identified in 44 tissues of the GTEx consortium^3,4^. We have adapted the Regulatory Trait Concordance (RTC) score^5^, to on the one hand measure the tissue sharing probabilities of eQTLs, and also to calculate the probability that a GWAS and an eQTL variant tag the same underlying functional effect. We show that our tissue sharing estimates significantly correlate with commonly used estimates of tissue sharing. By normalizing the GWAS-eQTL probabilities with the tissue sharing estimates of the eQTLs, we can estimate the tissues from which GWAS genetic causality arises. Our approach not only indicates the gene mediating individual GWAS signals, but also can highlight tissues where the genetic causality for an individual trait is manifested.

---

Over the last decade, Genome Wide Association Studies (GWAS) have become the norm in describing genetic variants associated with common complex human diseases and traits^1,6^. Although we have accumulated an impressive number of GWAS findings, the vast majority of the variants identified lie in the non-coding genome^7^, rendering their biological interpretation difficult. Furthermore, GWAS find genetic markers associated with organismal traits, and fail to pinpoint the specific tissues causing these associations^8^. Regulatory variants, like expression quantitative trait loci (eQTLs), identified in multiple tissues could aid greatly in the interpretation of GWAS results not only by linking the non-coding genome to genes, but also by identifying the causal tissues behind the genetic associations^2,9,10^. The Genotype Tissue Expression (GTEx) project was founded with the intention of characterizing eQTLs across multiple tissues^4^, and currently comprises 44 tissues from 449 individuals (70-361 samples per tissue) for a total of 7051 transcriptomes (**Supplementary Figure 1**). This makes GTEx the ideal dataset to estimate tissues from which the genetic causality of a GWAS trait arises. Here we aim to address this question by first, assessing the tissue sharing of eQTLs (the probability of an eQTL identified in one tissue being active in another tissue) on an individual variant basis, and then using these tissue sharing estimates to infer the tissues where GWAS variants exert their function.

For a given eQTL discovered in one tissue, we wanted to derive the probability that this eQTL is active in each of the other 43 tissues. We have previously described the Regulatory Trait Concordance (RTC) score, which tests whether co-localizing GWAS and eQTL variants (two variants that fall into the same genomic region delimited by recombination hotspots) are tagging the same functional variant^5^ (**Supplementary Methods** & **Supplementary Figure 3**). This method can easily be extended to assess tissue sharing between eQTLs identified in two separate tissues (**Supplementary Methods**). However, the RTC score is not a probability in itself and is affected by the number of variants and the linkage disequilibrium (LD) in a given region. Therefore, we derived a probability from the RTC score by simulating two scenarios for each region: (1) two variants tagging different functional effects (H0) and (2) two variants tagging the same functional effect (H1). Subsequently we generate a distribution centered on the real RTC found in the region and quantify the overlap between this distribution and simulated RTC scores under H0 and H1. We then apply the Bayes’ theorem, in conjunction with the overall tissue sharing estimates found by the π_1_ statistic^11^, to compute a probability of shared functional effect, which we call P(Shared), for a given RTC score in a given region (**Supplementary Methods, Supplementary Figure 4, Supplementary Figure 5 & Supplementary Figure 6**). By converting the RTC score into a probability we create a metric that accounts for the differential power of calling shared functional effects in different regions and which can be used in discovering tissue specificity of eQTLs.

Having a probability of sharing between two variants allows us to estimate tissue sharing of eQTLs amongst the 44 GTEx tissues. The gold standard methods used to quantify tissue sharing of eQTLs, such as the π_1_ method, estimate overall sharing, thus we aimed to estimate individual tissue sharing probabilities of each eQTL using π_1_ as a baseline. In order to ascertain a near-complete list of cis-eQTLs, we conducted a conditional cis-eQTL discovery and identify 858-13259 independent cis-eQTLs at FDR = 5% (**Supplementary Methods, Supplementary Figure 2**). Subsequently we took the union of eQTLs identified in all of the tissues (**Supplementary Methods**) and calculated sharing probabilities using the methodology described in the previous paragraph. We find a high degree of sharing amongst biologically related tissues. For example, brain tissues form a cluster with high sharing amongst themselves, coronary artery eQTLs are shared the most with aorta, and the uterus and ovaries share the most eQTLs (Figure 1a, **Supplementary Table 1**). We compared these tissue 66 sharing estimates to the more commonly used π_1_ estimates^11^ and find that the two metrics are significantly positively correlated (r = 0.933, p < 1e-300, Figure 1b), confirming the validity of our approach. The advantage of RTC over the π_1_ estimates is that RTC can assess the tissue sharing probabilities of an individual variant, whereas π_1_ estimates the overall sharing and cannot directly make a statement about individual eQTLs.

**Figure 1.**
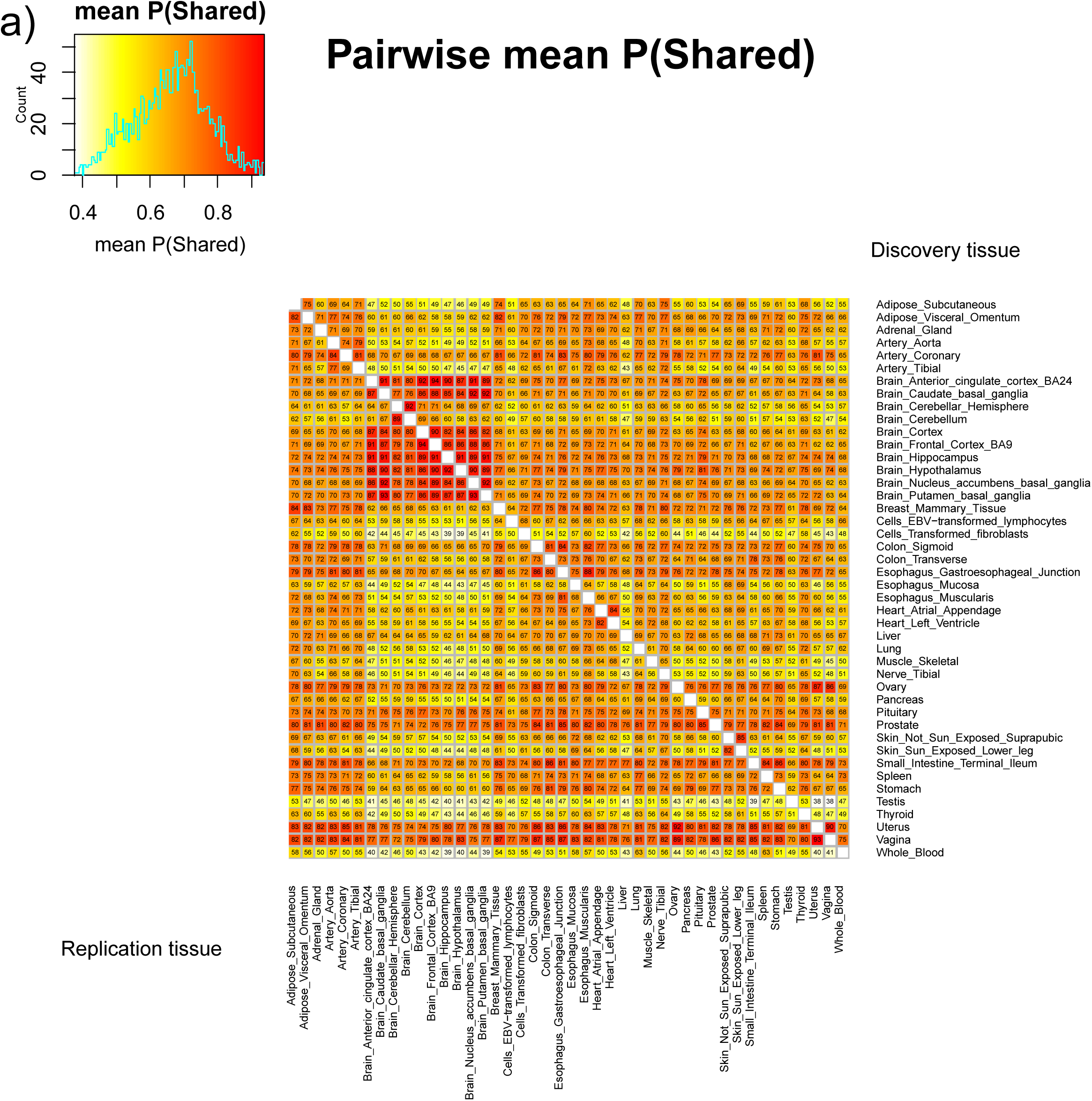
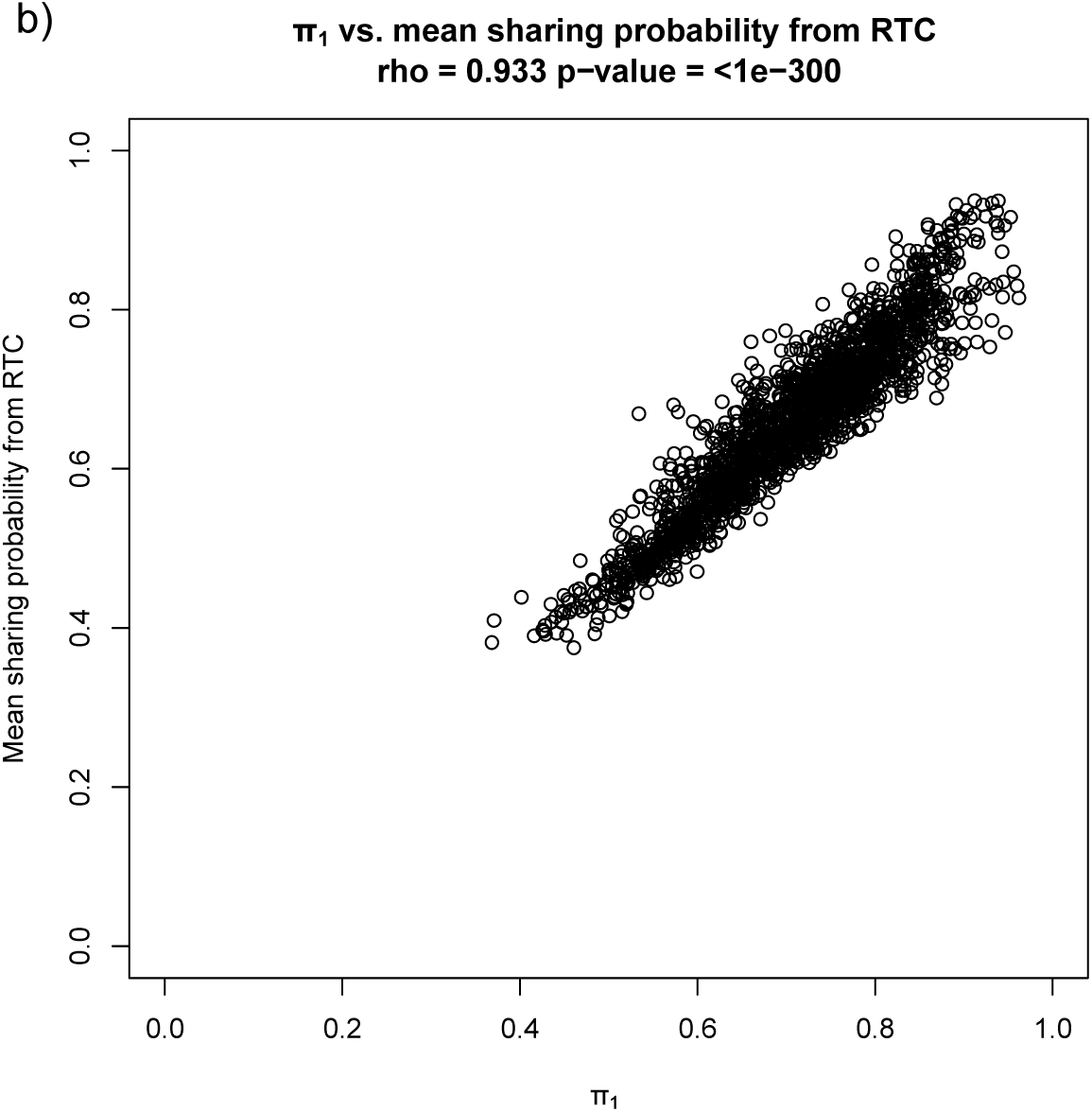
**(a)** Tissue sharing matrix based on sharing probabilities calculated through RTC. Rows correspond to the discovery tissue and columns to the replication tissue. Cells contain the mean probability multiplied by 100.**(b)** Correlation between the mean tissue sharing estimated from RTC and the π_1_ estimate, showing a significant positive correlation between these estimates.

Unlike the π_1_ estimates, our RTC-based probability of sharing can be used to find the most likely set of tissues where the eQTL effect is active. This is accomplished by enumerating the sharing probabilities of an eQTL in all combinations of the 44 tissues (**Supplementary Methods**). Moreover, we record the frequency of other tissues identified in the set of most likely tissues for an eQTL. The distribution of number of tissues an eQTL is likely active in show that the majority (94%) of the eQTLs are shared with at least one other tissue, in agreement with previous findings^4,12,13^ (Figure 2a). Furthermore, the number of tissues with shared effects decreases sharply as the number of tissues increases, but there is a slight enrichment for eQTLs active across most or all of the 44 tissues (Figure 2a). When we compare the eQTL sharing estimates among tissues for significant eQTLs found in each of the tissues, we discover two classes of tissues. Whereas the majority of the tissues exhibit higher degrees of tissue sharing, some tissues like testis and whole blood show a higher degree of tissue specificity (Figure 2b, c, e, **Supplementary Table 2**). Since each eQTL identified in a given tissue was predicted to be active in a set of other tissues, we next identified the most frequent other tissues included across all these sets. This was done to measure the global impact of the individual estimates, unlike the tissue sharing comparison in the previous section where we only quantify the global sharing between tissues. The results indicate for tissues with biologically meaningful similarity, shared eQTL effects are also more frequently observed. For example, brain tissues are most similar to other brain tissues, ovaries are most similar to uterus and vagina tissues, and heart 89 left ventricle is most similar to heart arterial appendage (Figure 2d & f, **Supplementary Figure 9**, **Supplementary Table 3**). In summary, our methodology uncovered two types of tissues, either with high degrees of tissue specificity or with high tissue sharing, and showed individual tissue sharing estimates identified biologically relevant tissues as shared, indicating the RTC method is capable of assessing tissue specificity on a variant by variant basis.

**Figure 2.**
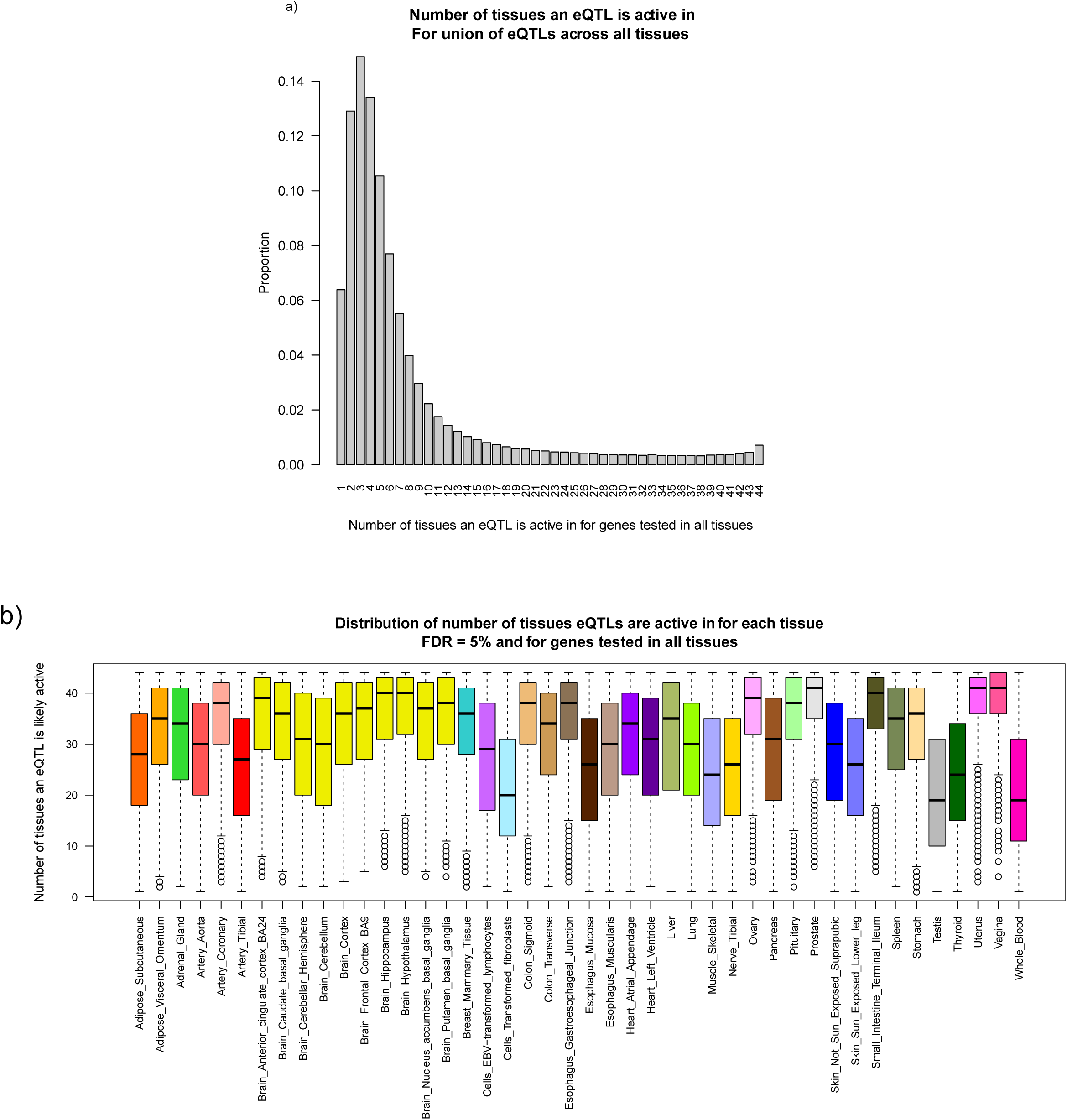
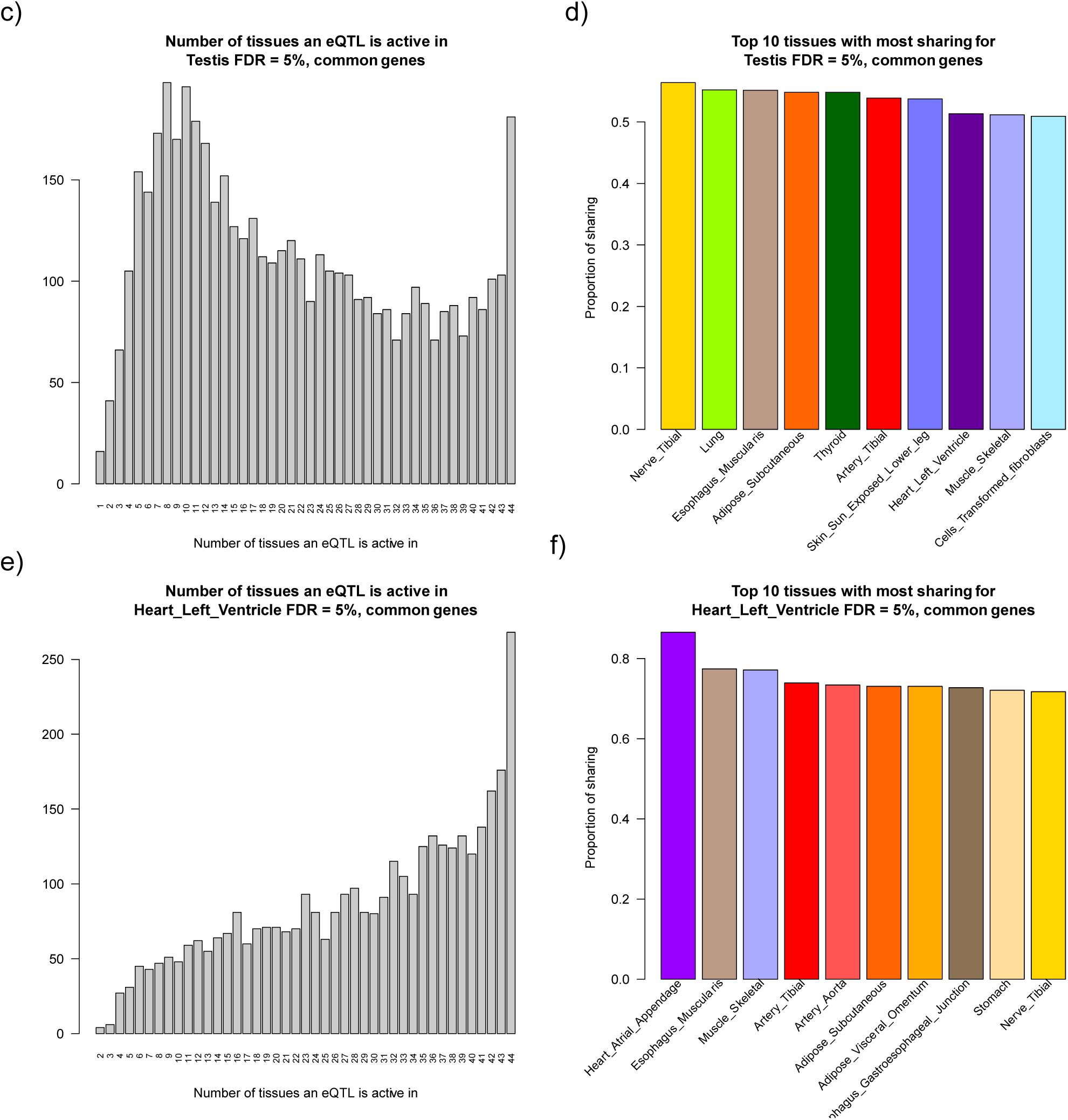
Frequency distribution of the number of tissues that an eQTL is activein (plotted for theunion of eQTLs across all tissues)which shows that most eQTLs are shared withat least one or a few other tissues, while eQTL sharing among high numbers of tissues is rare **(b)**Distribution of number of other tissues ane QTL is active in for the significant (FDR = 5%) eQTLs in each of the tissues. We see two types of tissues, one where the majority of eQTLs are shared with other tissues like the brain tissues and another set where there are higher rates of tissue specificity like test is and whole blood.**(c)**Test is as an example ofa tissue with a higher degree of tissue specificity of eQTLs and **(d)** the top 10 tissues that share eQTLs with test is.**(e)**Heart left ventricle as an example of a tissue that shares most of its eQTLs with other tissues and**(f)**the top 10 tissues that share eQTLs with left ventricle.

Given that GTEx comprises a wide range of tissues and our novel methodology can assess tissue sharing of each eQTL variant identified in these tissues, we are in an unprecedented position to infer candidate causal regulatory effects and their genes that may mediate the GWAS variants. Since RTC uses only the discovered GWAS variant, we are able to test GWAS-eQTL overlaps in all known GWAS variants, and were not limited to GWAS signals with available summary statistics or raw data, which unfortunately is very sparse. To this end we downloaded the NCBI GWAS catalogue^7^ and filtered the complete list of 15929 GWAS variants to include 5751 genome-wide significant associations (p < 5e-8) that overlapped with GTEx variants, and ran the RTC analysis with the independent significant eQTLs (FDR = 5%) from each of the tissues, resulting in 4664 GWAS variants that co-localized with eQTLs. We observe a large enrichment of high RTC scores across the GWAS-eQTL co-localizations confirming, as previously described^5,9,14^, that GWAS variants frequently manifest their effects through regulatory effects (Figure 3a). We also observe a bimodal distribution for probabilities of GWAS and eQTL tagging the same functional effect, where the majority of the probabilities are close to 0, but there also is an enrichment for high probabilities (Figure 3b). We have previously shown that RTC score is a better estimate of the causality between two variants than other pairwise LD metrics, r^2^ and D’^5^. When we compare the RTC score between two variants to their r^2^, we observe that high r^2^ generally means a high RTC score, however there many causal links found by RTC and missed when using r^2^ as a metric, extending our previous finding that RTC is preferable to r^2^ when predicting causality (Figure 3c, **Supplementary Figure 10, Supplementary Table 4**). Finally, we tested how the probability of sharing, as calculated with our new methodology varies with the raw RTC score and show that this probability behaves as expected with high RTC scores indicating a high probability of shared functional effects between the GWAS and eQTL variants. However, the probability is highly variable across regions that share the same RTC score, indicating the necessity of calculating this probability on a region by region basis (Figure 3d).

**Figure 3.**
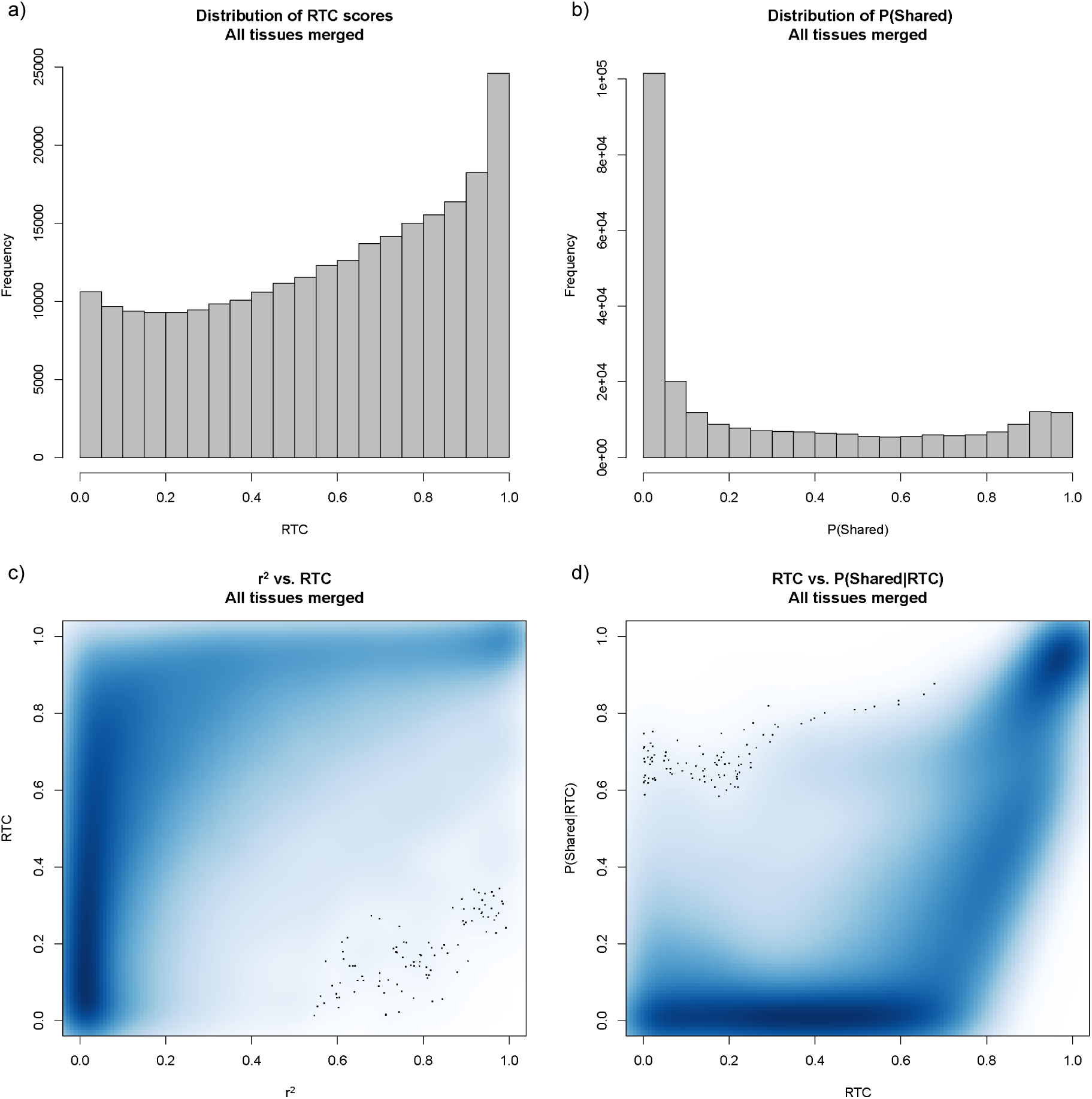
RTC score compared to other pairwise variant metrics. **(a)**Frequency distribution of RTC values for eQTLs from 44 tissues shows an enrichment in high RTC scores. **(b)** The distribution of the probabilities of GWAS and eQTL variants tagging the same functional effect. **(c)**RTC score compared to r^2^. High RTC tends to mean high LD between the two variants however low LD does not necessarily result in a low RTC, indicating that the RTC score is independent from the LD between the two variants.**(d)**Sharing probability calculated from simulations compared to the raw RTC score. Low RTC scores are much less likely to be shared than high RTC scores however thereis substantial variation between regions.

Although GWAS provide a list of markers that predispose to a certain disease or trait, they fail to identify the tissues where the genetic causality arises. Given that we can test all filtered GWAS signals for eQTL overlap, we can attempt to answer this question. In order to do so, we need to know not only whether co-localizing GWAS and eQTL variants are tagging the same functional effect, as inferred by RTC, but also the tissue-wide activity of the eQTL in question. We expect that weighing the probability of GWAS and eQTL variants being due to the same functional effect with the tissue sharing of the eQTL should increase our power in detecting the causal tissue behind the genetic associations of a GWAS trait. To do so, for each eQTL in a given tissue that co-localizes with a GWAS variant, we divide the probability of GWAS variant and eQTL tagging the same functional variant, with the sum of tissue sharing probabilities of that eQTL in that tissue. This enables us to weigh the GWAS-eQTL probabilities so that tissue specific eQTLs will contribute to a tissue’s GWAS enrichment more so than eQTLs that are shared with many other tissues. Next, for each disease in each tissue we divide the sum of the normalized GWAS-eQTL probabilities from the previous step with the number of independent eQTLs in the tissue, thereby controlling for the differential power of discovery amongst the 44 tissues, and this is defined as our enrichment metric. We show that by using our normalization technique we can significantly reduce (Mann-Whitney p = 1.5e-21, **Supplementary Figure 11**) the correlation between the number of eQTLs in the tissues and the GWAS enrichment metric, thus allowing us to estimate the relative contribution of tissues to the genetic causality of a trait.

We investigated the overall pattern in tissue causality of GWAS traits and looked at specific examples. Therefore, for each GWAS trait we rank our normalized enrichment metric for each of the tissues. Those tissues that are ranked higher are estimated to contribute more to the genetic causality of a GWAS trait. We discover that liver is the top tissue implicated in most of the GWAS traits (12%), which include, expectedly, variety of lipid measurements^15,16^ and uric acid levels^17^ (Figure 4a, **Supplementary Figure 12**, **Supplementary Table 5**). Brain tissues are the top tissues relating to traits like height^18^, schizophrenia^19,20^, and age of onset of puberty^21^. Furthermore, for traits where we have a biological prior of a causal tissue and where this tissue is assayed in GTEx, this tissue tends to be the most likely tissue discovered by our methodology. For example, the top causal tissue for coronary heart disease is coronary artery followed by liver; for schizophrenia the top tissues are brain tissues and for lipid metabolism traits like total cholesterol levels the top tissue tends to be liver (Figure 4b, c, d). Thus, we show that by having access to eQTLs from multiple tissues and controlling for the tissue specificity of eQTLs using our novel methodology, we can estimate the relevant tissues from which the genetic causality of GWAS traits arise.

**Figure 4.**
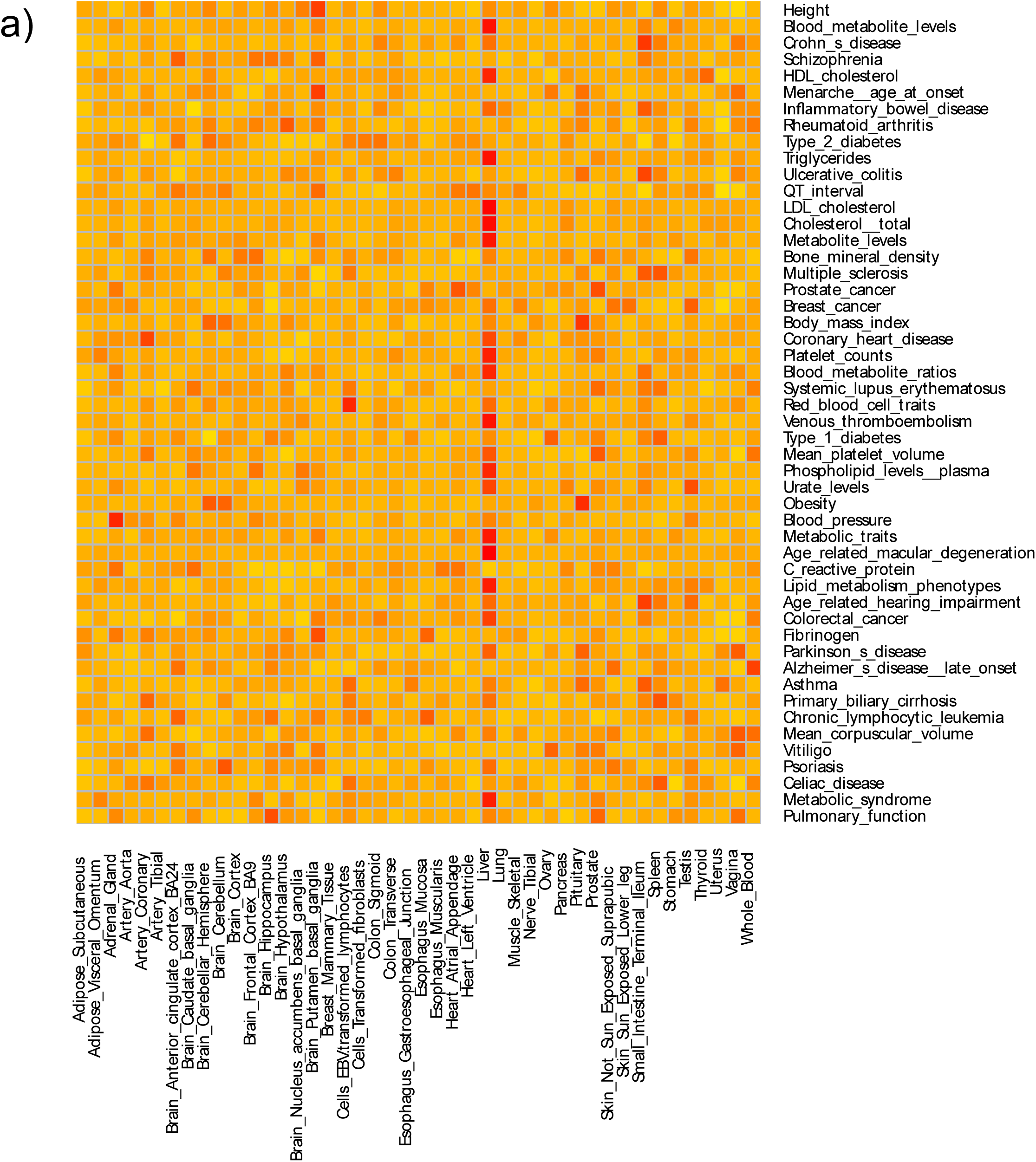
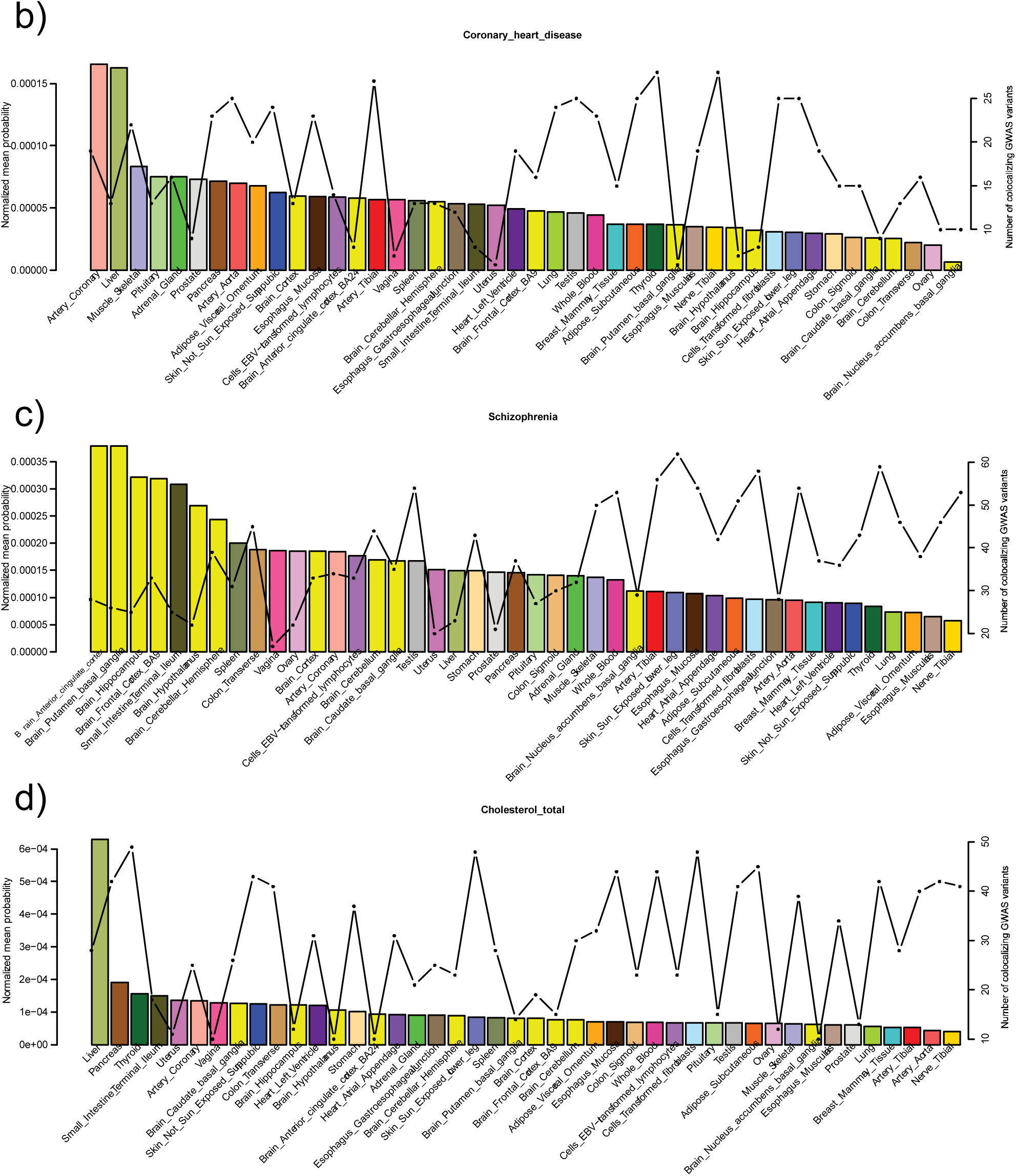
**(a)**Heat map of the relative likelihood of any tissue to be causally linked to any of the top 50 traits with the highest number of GWAS variants in the NCBI GWAS catalogue. Plotted are the relative normalized probabilities, which are centred and scaled per disease.The darker shades of red indicate higher likelihood of GWAS genetic causality is acting through this tissue. Rows correspond to the GWAS trait and columns to the tissues. Examples of traits with a prior on a biologically causal tissue:coronary heart disease **(b)**, schizophrenia **(c)**, total cholesterol **(d)**. On the primary y-axis the normalized probabilities per tissue are plotted as bars and onthe secondary y-axis number ofGWAS variants that co-localized with eQTLs per tissue are plotted as a line.

Since we estimated the tissue causality profiles for GWAS traits, we can compare the causal genes for the GWAS associations between tissues likely contributing to the genetic causality of GWAS traits and those that are not. We examined the rs12740374 variant in the 1p13 locus, which is not only associated with coronary artery disease^22,23^ and lipid measurements^24^, but is also one of the few GWAS non-coding loci where the mechanistic causes are well established^25^. Liver is a key tissue in both heart disease and lipid measurements (Figure 4b, d), and in liver the causal gene for the rs12740374 association is correctly identified as *SORT1*^25^, P(Shared) = 1. In tissues that do not contribute highly to the genetic causality of these traits, like testis and whole blood, we incorrectly identified another nearby gene, *PSRC1*, as the putative causal gene, P(Shared) = 0.96 and 0.97, respectively (Figure 5, **Supplementary Table 6**). Importantly, the tissues where *SORT1* is correctly identified contribute significantly (Mann-Whitney p = 0.0007) more to the genetic causality of heart disease and lipid levels than tissues where the causal gene is different (**Supplementary Figure 13**). This result shows the importance of identifying the causal tissues for GWAS traits, before stating which genes may be responsible for these associations.

**Figure 5.**
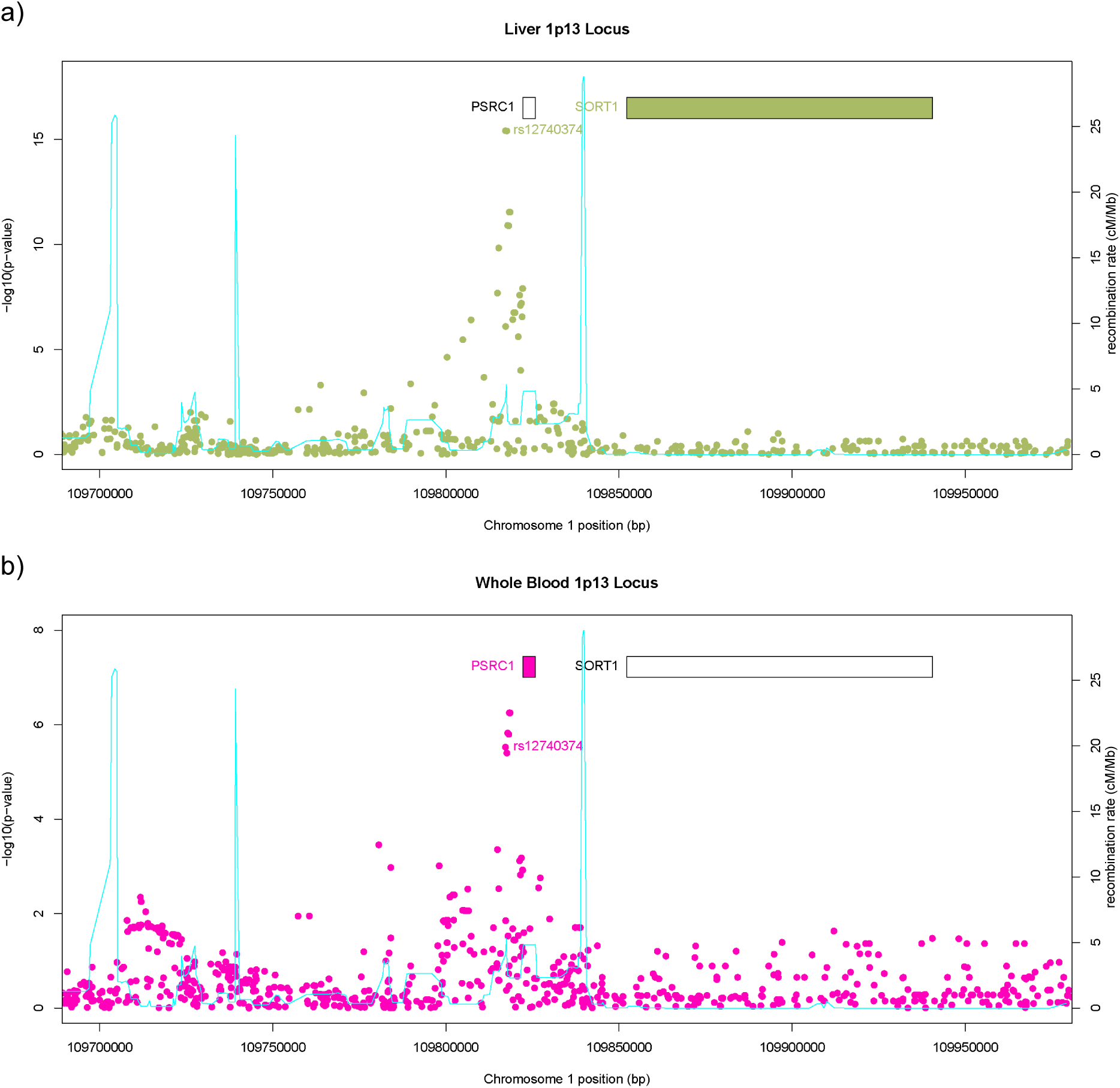
The coronary artery disease (CAD) and lipid levels associated GWAS 1p13 locus eQTLs in liver**(a)** and whole blood**(b)**. Points are the -log10(p-value) eQTL associations for *SORT1* in liver and *PSRC1* in whole blood. The cyan line is the recombination rate, given in the secondary y-axis, and the boxes highlight the positions of the two genes. In both tissues the best eQTL association is genome-wide significant (FDR = 5%), however the eQTL gene, for which the eQTL and the causal rs12740374 variant are tagging the same functional effect as identified by our method, is different. Liver, which we estimate to play a key role in the development of both CAD and regulation of lipid levels, correctly identifies the*SORT1* as the causal gene for this GWAS association, however the more easily collectable whole blood tissue, which is estimated not to contribute to these traits, fails to do so. If we had just whole blood eQTLs, and did not know the tissue causality profile for these traits, we would have incorrectly identified *PSRC1* as a putative causal gene.

Finally, we asked how different diseases with shared pathophysiology differ with respect to which tissues contribute to their genetic causality. To this end we investigated autoimmune and cardiometabolic diseases and used hierarchical clustering to group the individual diseases according to their relative tissue causality profiles. Among the autoimmune diseases we find that Crohn’s disease and ulcerative colitis form a cluster, whereas celiac disease has a different tissue causality profile. Type 1 diabetes and lupus seem most similar to each other and rheumatoid arthritis, vitiligo, and psoriasis appear markedly different when compared to other autoimmune disorders (Figure 6a). For cardiometabolic diseases, blood pressure related traits, coronary heart disease and type 2 diabetes form a cluster while stroke, where a strong effect in the brain is observed, is the outlier in these types of disorders (Figure 6b). We demonstrate that by comparing the tissue causality profiles of GWAS diseases we can begin to disentangle the common as well as diverging biology underlying their development.

**Figure 6.**
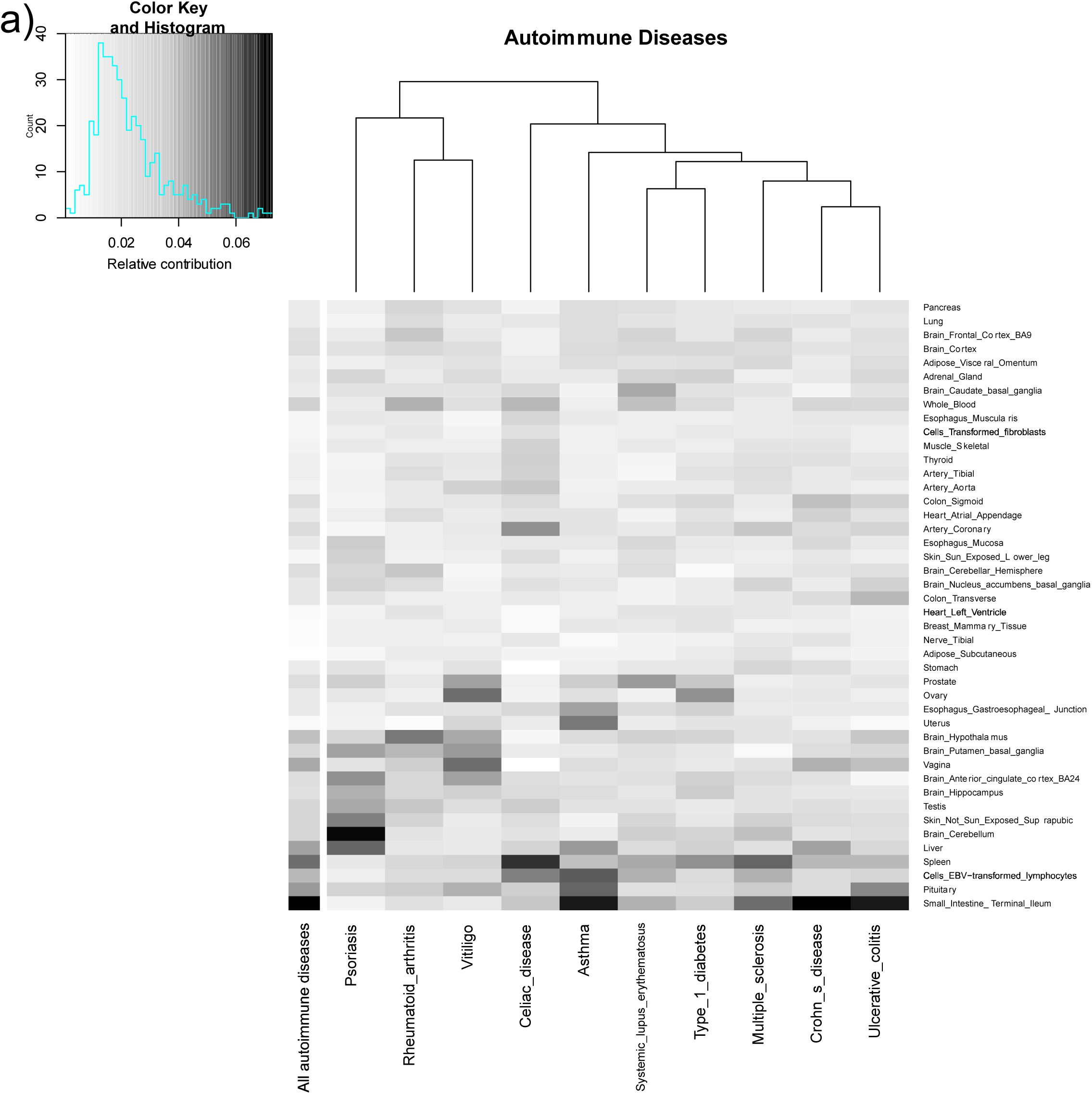
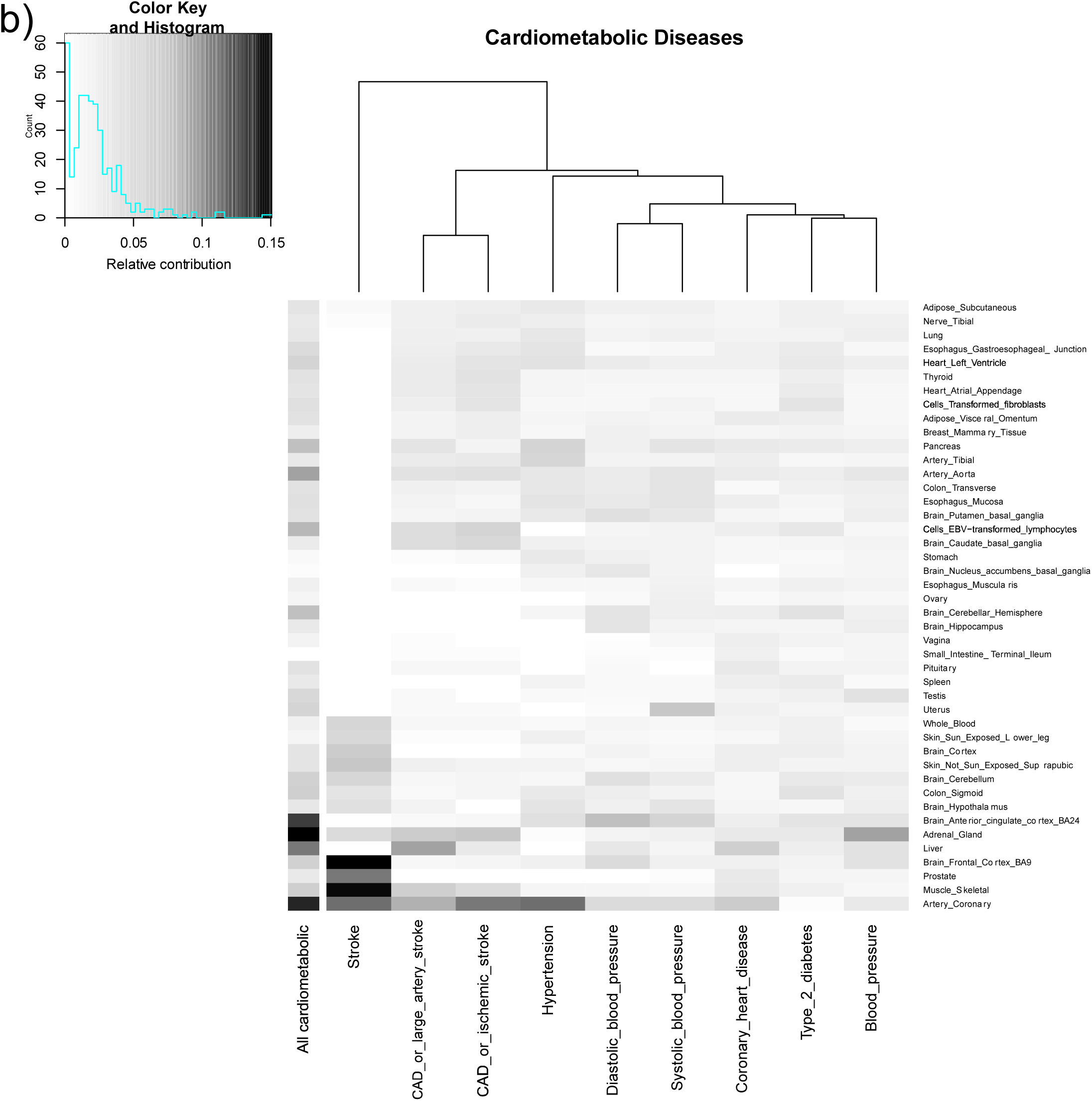
Relative contribution of tissues to the genetic causality of autoimmune diseases **(a)** and cardiometabolic disorders**(b)**. Rows list tissues and columns list diseases. Darker shades represent higher contribution per tissue. The left most column shows the relative tissue contributions across all diseases combined. The hierarchical clustering of the diseases is shown as a dendrogram.

Here we describe a novel approach that is designed to estimate the causal tissues underlying the genetic causality of GWAS traits, using eQTLs identified by the GTEx consortium. Given the tissue and sample size limitations, there is still room for improvement in determining true tissue causality profiles for GWAS traits. However, our analysis represents an unbiased and complete profiling of the 181 tissue contributions to GWAS genetic causality in an unprecedented scale. As the sample sizes and the number of tissues assessed for eQTLs, and our resolution of the genetic etiology of complex disorders increase, we expect our methodology to yield even more powerful conclusions. We believe this type of approach will be paramount in the interpretation of new GWAS results using a publically available dataset, like GTEx, and will aid in the design of downstream functional experiments to identify the mechanistic causes of complex disorders and traits, as well as new avenues of treatment and prevention.

## Data access

The data used in this paper is available for controlled access at dbGaP (accession: phs000424.v6.p1).

## Methods

are described in the supplementary methods and figures file available in the online version of this paper.

## Acknowledgements

This research was support by grants from NIH-NIHM (GTEx), European Commission FP7, European Research Council, and Swiss National Science Foundation. The computations were performed at the Vital-IT Centre for high performance computing of the SIB Swiss Institute of Bioinformatics.

## URLs

GTEx dbGaP http://www.ncbi.nlm.nih.gov/projects/gap/cgi-bin/study.cgi?study_id=phs000424.v6.p1

GTEx Portal http://gtexportal.org/home/

QTLtools https://qtltools.github.io/qtltools/

Vital-IT http://www.vital-it.ch/

## Author contributions

H.O. and E.T.D designed the study. H.O., A.A.B., and O.D. conducted the 202 analysis and developed software. A.C.N designed the original RTC method. N.P. tested the software. H.O. wrote and E.T.D edited the paper. GTEx Consortium generated the data.

